# Neuroimaging of intimate partner violence can underrepresent the most affected survivors

**DOI:** 10.64898/2026.09.23.753381

**Authors:** Craig W. McFarland, Aaron Kucyi, Eve M. Valera

## Abstract

Neuroimaging is becoming an increasingly prominent tool to understand intimate partner violence (IPV) and partner-inflicted brain injury (BI). Yet, by nature of IPV and partner-inflicted BI, those most affected may be the least likely to be eligible, participate, and produce usable data in neuroimaging studies. A critical question therefore remains of who is represented in neuroimaging research on IPV and related BI. Here, we examined representativeness at three major stages of neuroimaging sample construction using screening records from 352 women and a subsequent cohort of 207 enrolled women with lifetime exposure to physical IPV. At the eligibility stage, three in five women eligible for behavioral research were deemed ineligible for magnetic resonance imaging (MRI) at initial screening. At the participation stage, the 61 women who underwent resting-state functional MRI (fMRI) had more education, were more likely to be employed, and reported lower psychiatric and neurobehavioral symptom burden than enrolled women who did not undergo neuroimaging. Finally, at the quality-control stage following fMRI, motion-based exclusion disproportionately removed women who had sustained the most partner-inflicted BIs. Our findings empirically reveal that eligibility, participation, and quality control each bias the demographic, clinical, abuse, and BI burden represented in an fMRI sample, and therefore have key implications for generalizability. Measuring and reporting representation bias can help identify whose experiences are underrepresented in the evidence, and accordingly reduce the risk that IPV neuroimaging research understates the very harms it seeks to characterize.

## Introduction

Intimate partner violence (IPV) is widespread, with an estimated one in three women experiencing IPV [1]. Blows to the head, strangulation, and other forms of neurotrauma are common, and brain injury (BI) is a frequent consequence [2]. Partner-inflicted BI is often repetitive [3] and associated with cognitive impairment [4], debilitating neurobehavioral symptoms [5], and psychiatric difficulties [3]. Despite this burden, partner-inflicted BI often remains “invisible” [6]: injuries are rarely documented when they occur, their symptoms are often attributed to psychiatric distress, and the underlying BI may go unrecognized by survivors and medical professionals, leaving many without appropriate care [2,5].

To address this “invisible” burden, neuroimaging has emerged as a tool for characterizing the neural consequences of IPV and associated BI. Complementing behavioral evidence, neuroimaging studies have identified differences in cortical morphometry [7], white matter microstructure [8], anterior cingulate neurochemistry [9], structural connectivity [10], and functional connectivity [6].

Neuroimaging approaches, however, are particularly demanding for participants and investigators. Eligibility depends on scanner safety and clinical criteria, while participation requires time, transportation, scheduling flexibility, and tolerance of lengthy procedures and the scanner environment. Population-based and perioperative studies show that imaging participants differ from nonparticipants in education [11,12], physical and psychological health [13–15], cognition, and clinical status [16]. These differences can alter estimated brain-behavior associations [11,17–19]. Selection continues after acquisition because motion and quality-control exclusions vary with factors such as cognition [20], autism severity [21], physical and psychological health [22], sociodemographic status [23], and the quality-control pathway applied [24].

The concerns about representativeness identified in the broader neuroimaging literature may be especially consequential in IPV research. Women exposed to IPV disproportionately come from racial and ethnic groups that remain underrepresented in neuroimaging [25] and from socioeconomically disadvantaged populations [5]. Ongoing abuse may restrict safe communication, travel, scheduling, and time away from caregiving [26,27], while pervasive stigma may constrain disclosure and health-seeking behavior [5,28–30]. Among women who have sustained one or more BIs, associated cognitive and neurobehavioral impairments may compound barriers to completing lengthy and procedurally demanding studies [3,4]. Factors shown to bias quality-control exclusions, including cognition [20] and physical health [22], are also prevalent in IPV. Together, these overlapping pressures could produce a final analytic sample that underrepresents women affected by the greatest social, clinical, and injury-related burden.

As neuroimaging becomes increasingly prominent in IPV research, especially with the high prevalence of partner-inflicted BIs, a critical gap is understanding how imaging eligibility, participation, and quality control shape who is represented in the resulting evidence. To address this gap, we examined three major stages of neuroimaging sample construction using screening records from 352 women and an analytic cohort of 207 enrolled women with lifetime exposure to physical IPV who completed two structured interviews to assess BI burden. First, at the eligibility stage, we determined which behaviorally eligible women also met neuroimaging criteria and how changes in those criteria affected projected access. Next, at the participation stage, we compared enrolled women who underwent fMRI with those who did not. Finally, at the quality-control stage, we tested whether motion-based exclusion within the fMRI sample varied with cumulative partner-inflicted BI. Separating these stages allowed us to identify when sample composition changed and which population was represented in the final fMRI analysis. Ultimately, our aim was to identify sources of selection and potential representation bias across IPV neuroimaging and inform strategies to improve scientific rigor and representation in this emerging field.

## Methods

### Participants

Data were obtained from a community cohort of women with lifetime exposure to physical IPV. Data were collected between 2020 and 2026 as part of an ongoing study of IPV-related BI [6,8]. Participants were recruited through community partners, an institutional online research platform, and social media. The analytic cohort included 207 enrolled women who completed the two structured BI interviews. A personal history of BI was not required for enrollment. The analyses examined imaging eligibility, imaging participation, and motion-based exclusion within this study.

Eligibility was assessed by telephone and recorded separately for remote behavioral participation and in-person imaging. Behavioral eligibility required an age of at least 18 years, at least one physically abusive incident by an intimate partner, capacity to consent, and sufficient English proficiency. Additional imaging exclusions included magnetic-resonance contraindications, pregnancy, a lifetime history of schizophrenia, bipolar disorder, or autism, moderate or severe traumatic BI, BI within the preceding three months, moderate or severe substance use disorder within six months, another neurological disorder that could affect study outcomes, and geographic distance that made travel to the imaging center impractical. These criteria did not affect behavioral eligibility.

It should be noted that, following difficulty with recruitment, three imaging criteria were changed early in the study. Current psychotropic medication use was removed, as was a criterion that excluded women reporting only one or two partner-inflicted BIs, and an upper age limit of 55 years was discontinued. The eligibility analysis includes decisions made under different criterion sets. We therefore classified each ineligible woman by whether she failed at least one exclusion criterion that remained in use or only failed criteria that were later removed.

The Mass General Brigham Institutional Review Board approved all procedures. The study was conducted in accordance with the Declaration of Helsinki, and all participants provided written informed consent.

### Brain injury burden

BI burden was assessed with two structured interviews administered by a trained examiner. The Brain Injury Severity Assessment [31] applies American Congress of Rehabilitation Medicine criteria [32] to events inflicted by an intimate partner. An event was classified as a BI when it involved traumatically induced loss of consciousness, peri-event amnesia, altered mental state, or focal neurological signs. The Ohio State University TBI Identification Method assesses lifetime injuries from any cause [33]. Events meeting criteria on either interview contributed to the count. Consistent with earlier reports from this cohort, altered consciousness attributed to anoxia or hypoxia during strangulation was also classified as a BI.

The primary exposure was cumulative partner-inflicted BI count. The BI distribution was strongly right-skewed, with a median of 1, IQR of 0–4, 90th percentile of 14, maximum BI count of 1,095, mean of 14.4, *SD* of 84.2, and adjusted Fisher-Pearson *G_1_* = 11.0. When women reported extraordinary levels of partner-inflicted BI, they estimated a BI frequency over a set duration. To illustrate, a woman sustaining two BIs a week across a ten-year abusive relationship would be calculated to be over a thousand events. To mitigate the influence of these estimated totals, we winsorized BI counts at 25, following prior work in this cohort [4,34]. Eleven of 61 women with rsfMRI data had BI counts over this cap. In addition, we log-transformed the count when examining its correlation with head motion.

We summarized BI count using the median and IQR, and quantified group differences with Cliff’s delta. For comparison with other measures, we report Hedges’ *g* on the winsorized count. The threshold of ten or more BIs used in the motion analysis was specified before this analysis on the basis of earlier work in the cohort [34].

### Symptom, socioeconomic and demographic measures

Depression was measured with the PHQ-9 [35], anxiety with the GAD-7 [36], and post-traumatic stress with the PCL-5 [37]. We summed standardized PHQ-9, GAD-7, and PCL-5 scores to create a combined symptom score and defined the upper quartile as the group with the greatest psychiatric symptom burden. Post-concussive symptoms were assessed with the 22-item Neurobehavioral Symptom Inventory (NSI) [38]. Total scores were calculated when at least 18 items were available. Following the established four-factor structure, items were grouped into vestibular (items 1–3), somatosensory (items 4–10), cognitive (items 11–14), and affective (items 15–22) subdomains, each scored as the item mean. Education was recorded in years, and current employment was recorded at interview. Participants self-reported race and ethnicity. Records without a selected race category were treated as missing. Childhood maltreatment was measured with the Childhood Trauma Questionnaire–Short Form [39]. Recent abuse severity was measured with the Composite Abuse Scale (Revised)–Short Form [40]. Ongoing abuse was recorded during the telephone screening, and time since the most recent partner-inflicted BI was recorded in years.

### Image acquisition and estimation of head motion

Resting-state fMRI was acquired with a 3-Tesla Siemens scanner at the Athinoula A. Martinos Center for Biomedical Imaging. Participants were instructed to lie quietly with their eyes focused on a crosshair projected onto the middle of a screen and to try to stay awake. Head motion was summarized as mean framewise displacement (FD) across four resting-state runs per participant, each lasting 6.5 minutes. We computed FD with the MCFLIRT tool in the FMRIB Software Library [41,42].

We examined mean FD thresholds of 0.15, 0.20, and 0.30 mm and designated 0.20 mm as primary. No single participant-level threshold is standard across studies. The ABCD processing pipeline uses 0.20 mm to censor individual frames and requires at least 375 retained frames for participant inclusion [43]. Power and colleagues censored frames above 0.50 mm [44], and Cosgrove and colleagues excluded participants with mean FD above 0.15 mm [20]. Analysis across three participant-level thresholds assessed sensitivity to this choice.

### Definition of the imaging samples

We defined the primary imaging group as women with rsfMRI data available before motion-based exclusion who had also completed prior behavioural assessments. Sixty-one of 207 enrolled women met this definition, so participation comparisons included 61 women with rsfMRI data and 146 without.

### Statistical analysis

For continuous characteristics, we report Hedges *g* as the bias-corrected standardized mean difference between women with and without imaging data. Percentile bootstrap 95% confidence intervals were calculated using 5,000 resamples. Mann-Whitney *U* tests were also calculated because several measures had skewed distributions. For strongly skewed count variables, we emphasized medians, IQRs, and Cliff’s delta. Binary characteristics are reported as odds ratios with the Haldane-Anscombe correction and 95% confidence intervals. Two-sided Fisher exact tests were used because several cells contained small counts.

We interpreted estimates according to their magnitude and precision rather than a fixed significance threshold [45,46]. We did not adjust for multiple comparisons because the analyses were descriptive and the estimates addressed distinct characteristics of one cohort. Analyses used Python 3.11 with pandas, NumPy, and SciPy.

### Missing data

Missing values were retained as missing, and no imputation was performed. Missing responses were not recoded as negative responses. The analyzed sample size is reported for each estimate. Time since the most recent partner-inflicted BI was available for 122 of 207 women. Among 211 women deemed ineligible for imaging, the magnetic-resonance contraindication item was assessed in 153. For each exclusion reason, we report the percentage of all ineligible women and the percentage of women assessed for that item.

## Results

### Eligibility, participation, and quality control shaped sample composition

We began by quantifying sample composition across three major stages of IPV neuroimaging (Fig. 1). Among 352 women screened as eligible for behavioral participation and given an imaging decision, 141 (40.1%) also met imaging criteria. Among the enrolled cohort of 207 women who completed two structured BI interviews, 61 (29.5%) had rsfMRI data. Among those 61 women, 55 (90.2%) met the primary head-motion quality-control criterion.

**Fig. 1.**
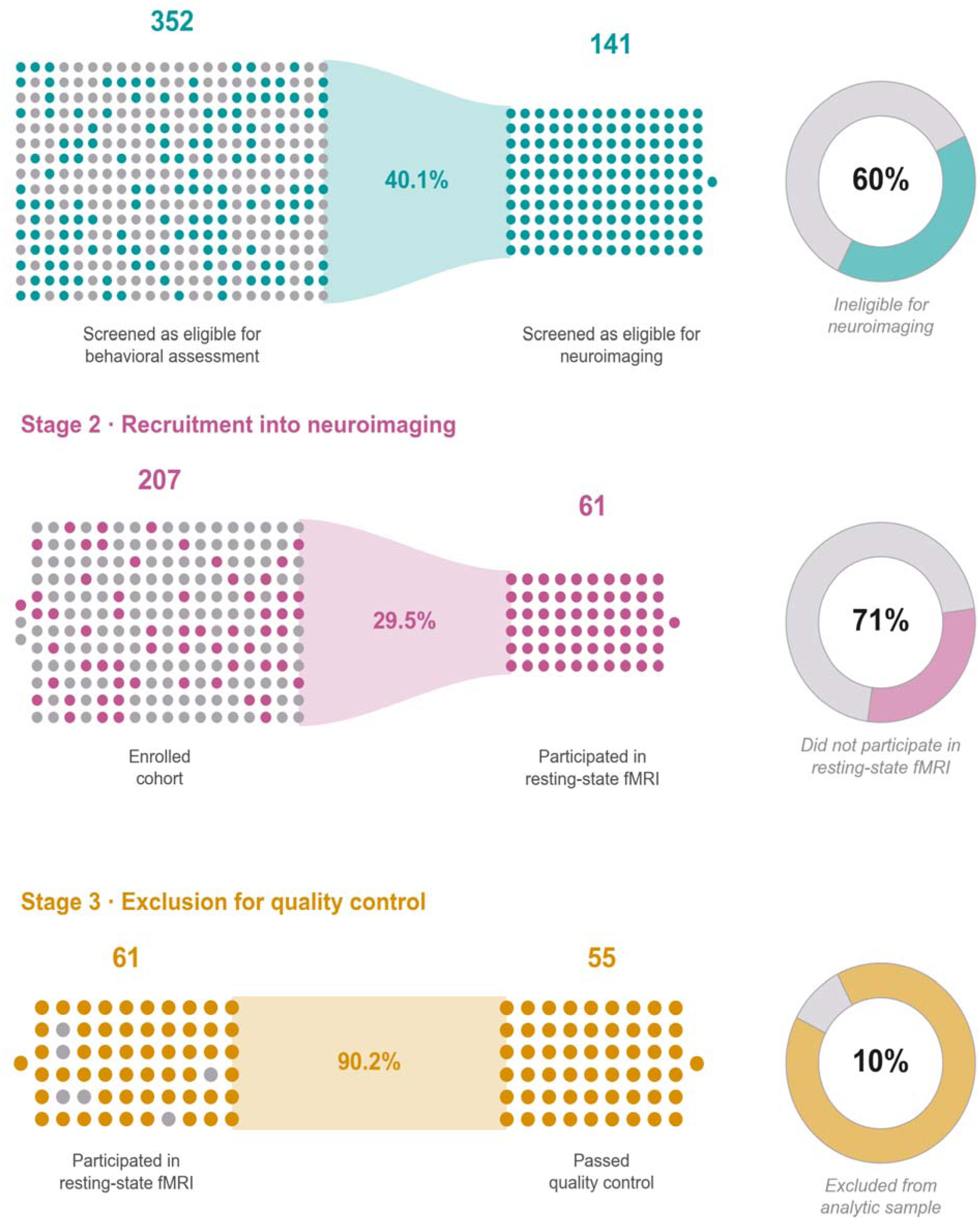
Data exclusion in the neuroimaging of IPV by selection stage. Each circle is one woman: the block on the left is everyone entering the stage, with those lost shown in gray among them; the band gives the percentage continuing; the block on the right is those who continue; and the ring gives the percentage lost. Of 352 women screened as eligible for behavioral assessment, 141 (40.1%) were eligible for neuroimaging. Of 207 enrolled women, 61 (29.5%) had rsfMRI data. Of those 61, 55 (90.2%) passed head-motion quality control at a mean framewise displacement below 0.20 mm. The stages do not share a denominator, and panels are scaled to a similar area, so counts should be compared within a panel rather than across them.

### Imaging criteria substantially narrowed behavioral eligibility

At the eligibility stage, neuroimaging studies of IPV span structural MRI [7], diffusion MRI [8], spectroscopy [9], and functional MRI [6], with screening criteria varying across experimental designs. To quantify how imaging screening altered who could enter our study, we compared behavioral and imaging eligibility decisions among 352 women screened as eligible for behavioral participation. Of these women, 141 (40.1%) also met imaging criteria, whereas 211 (59.9%) did not. Imaging criteria therefore excluded three in five women who could otherwise participate in behavioral research.

To identify the main sources of imaging ineligibility, we summarized every recorded reason among the 211 excluded women (Fig. 2 and Supplementary Table 2). Criteria used throughout the study included geographic distance from the imaging center in 64 women, a magnetic-resonance contraindication such as metal in the body in 58, a history of schizophrenia, bipolar disorder, or autism in 34, moderate or severe traumatic BI in 30, another neurological disorder in 29, a moderate or severe substance use disorder within the preceding six months or a chronic substance use history judged likely to confound results in 25, BI within the preceding three months in 17, and pregnancy in 1. The magnetic-resonance contraindication accounted for 27.5% of excluded women and 37.9% of the 153 women assessed on this item. Earlier recruitment phases also excluded women taking psychotropic medication, women reporting only one or two BIs, and women older than the study age limit. Psychotropic medication use was recorded for 105 women, representing 49.8% of those deemed initially ineligible.

**Fig. 2.**
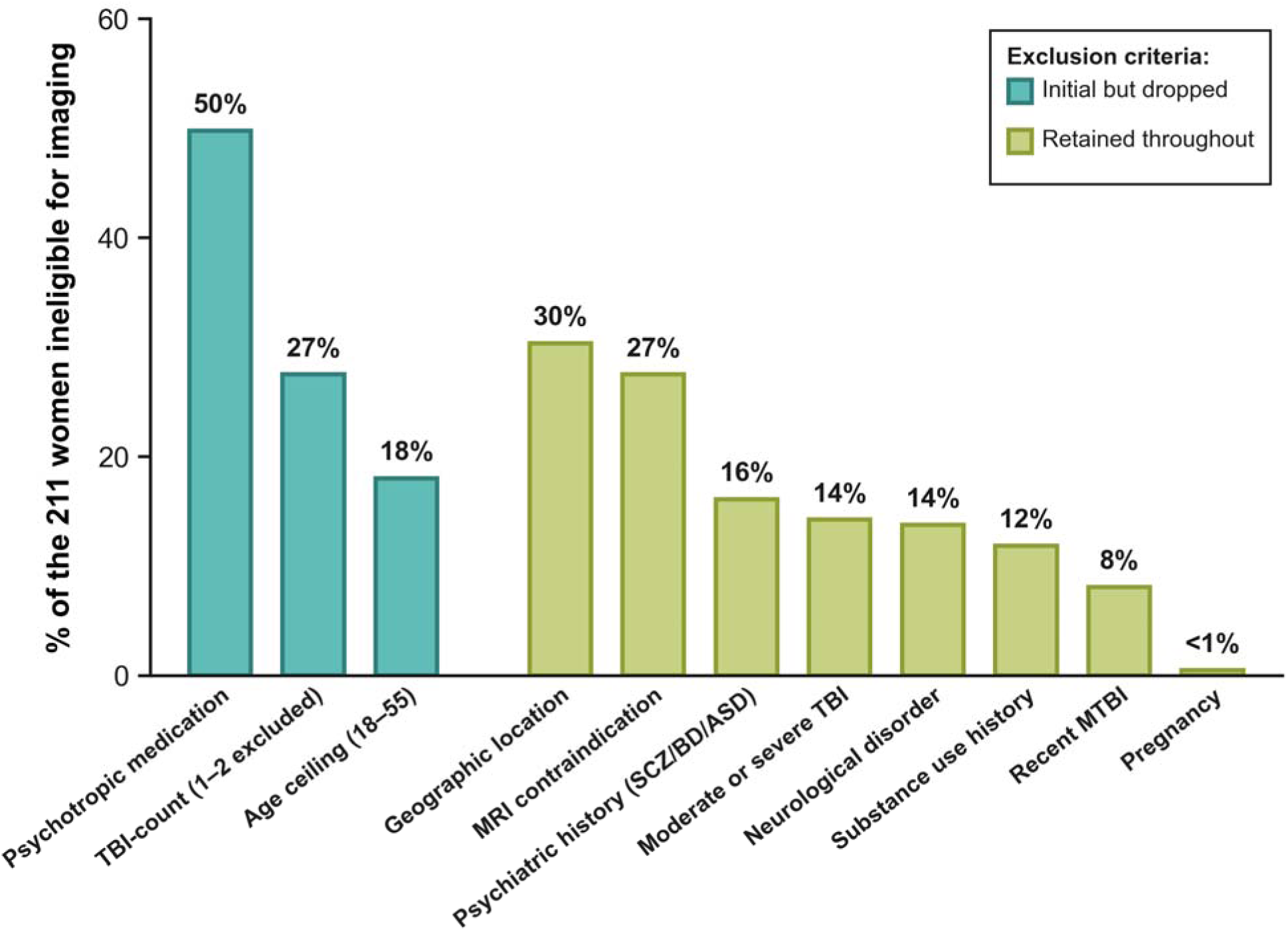
Breakdown of neuroimaging ineligibility. Percentage of the 211 behaviorally eligible women found ineligible for neuroimaging who met each criterion, grouped by whether the criterion has since been relaxed or dropped (teal) or was applied throughout (green). A woman could meet more than one criterion, so percentages sum to more than 100, and none of these criteria affected behavioral eligibility. The magnetic-resonance contraindication item was assessed in 153 of the 211, among whom its prevalence was 37.9% (Supplementary Table 2). Notably, the remaining 58 of the 211 were already excluded prior to being screened for contraindication. Forty-nine women (23%) failed only initial exclusion criteria that have since been relaxed or dropped.

To determine how changes in eligibility criteria affected access to imaging, we classified each ineligible woman by the criteria she failed. Psychotropic medication use had been removed, as had the criterion excluding women with only one or two partner-inflicted BIs, and the upper age limit of 55 years had been discontinued. Of the 211 women deemed ineligible, 161 (76%) failed at least one criterion that remained in use, and 49 (23%) failed only criteria that were later removed. Notably, fourteen (29%) of these 49 women subsequently underwent imaging after the criteria were relaxed.

### Imaging participants differed in social and clinical burden

At the participation stage, behavioral assessment and neuroimaging impose different demands, and prior studies show that imaging participants may differ systematically from nonparticipants [11–16]. To determine whether the women represented in the imaging sample differed from the broader enrolled cohort, we compared 61 women with rsfMRI data and 146 without these data. This comparison characterized differences between the behavioral and imaging samples before motion quality control. Women with imaging data had more education (16.6 versus 15.4 years, *g* = 0.42, 95% CI [0.13, 0.72]) and were more often employed (75% versus 59%, OR = 2.06, 95% CI [1.06, 4.00]). They reported lower depression (PHQ-9 8.1 versus 10.2, *g* = −0.37, 95% CI [−0.64, −0.09]), anxiety (GAD-7 6.9 versus 8.9, *g* = −0.35, 95% CI [−0.64, −0.07]), and post-traumatic stress (PCL-5 29.2 versus 35.3, *g* = −0.32, 95% CI [−0.59, −0.02]). Total neurobehavioral symptoms were also lower (NSI 20.3 versus 27.4, *g* = −0.50, 95% CI [−0.78, −0.21]), with standardized differences from −0.32 to −0.46 across the four NSI subdomains (Fig. 3 and Supplementary Table 1).

**Fig. 3.**
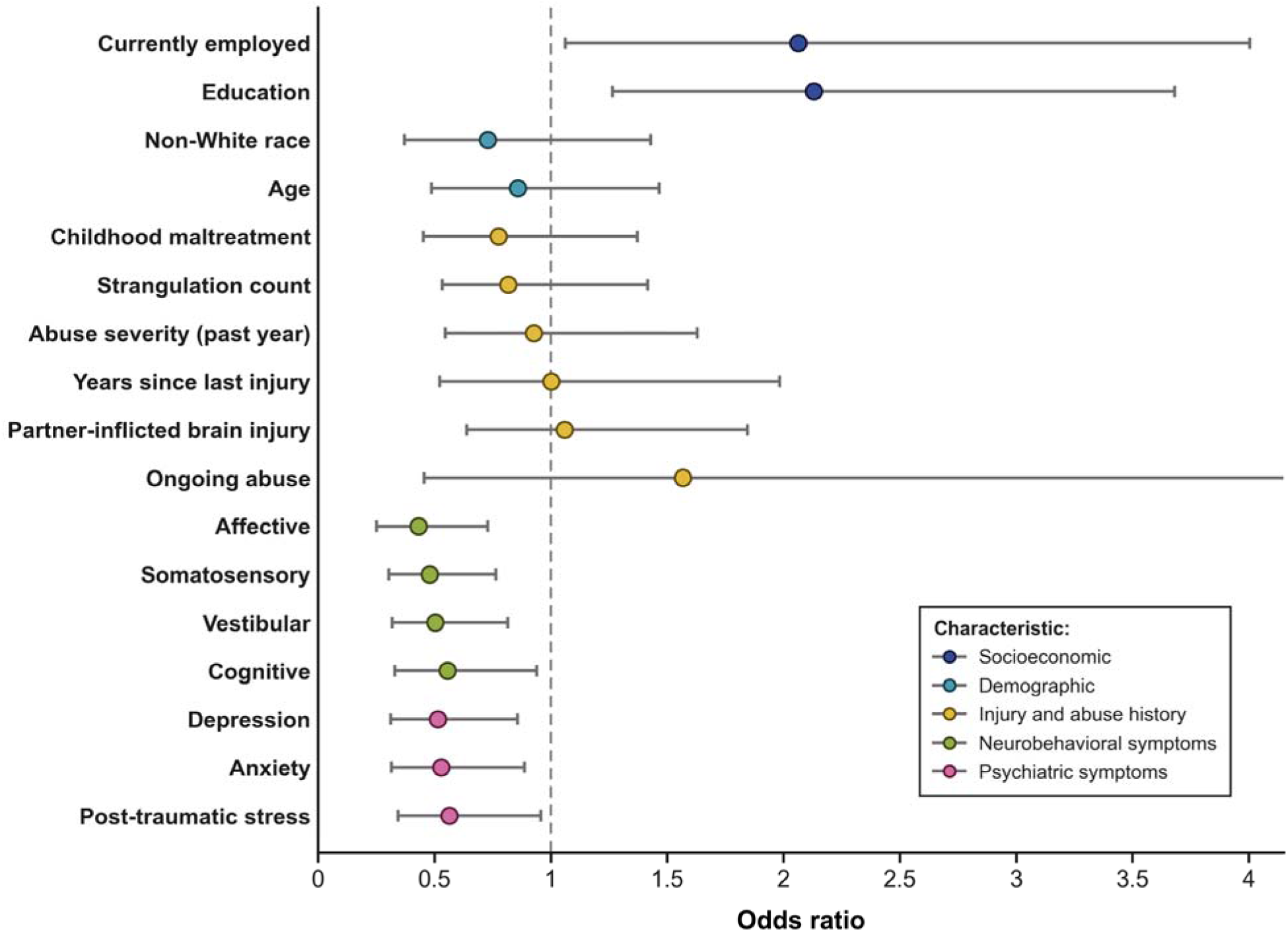
Representation bias in participation of neuroimaging study. Odds of having resting-state fMRI data for the 61 women who participated in neuroimaging against the 146 who did not. Points are odds ratios, bars are 95% confidence intervals, and color gives the domain. For current employment, non-White race, and ongoing abuse, estimates are odds ratios with 95% confidence intervals calculated using the Haldane– Anscombe correction. For the fourteen continuous characteristics, they are derived from Hedges *g* by ln(OR) = *d* × π/√3 [47,48]. Depression, anxiety and post-traumatic stress are assessed via the PHQ-9, GAD-7 and PCL-5, respectively. The neurobehavioral rows are domains assessed via the Neurobehavioral Symptom Inventory. BI and strangulation counts are winsorized at 25. Native estimates are in Supplementary Table 1. Estimates are interpreted by magnitude and precision [45,46].

To assess whether the behavioral-only and imaging samples differed by injury and abuse history and by demographic characteristics, we compared cumulative BI, strangulation-related BI, childhood maltreatment, past-year abuse severity, race, and age. Median cumulative partner-inflicted BI count was 1 [IQR 0–5] among women with imaging data and 1 [IQR 0–4] among those without (Cliff’s δ = 0.03). The winsorized standardized difference was *g* = 0.03 (95% CI [−0.25, 0.34]). Estimates were also small for strangulation-related BI count (Cliff’s δ = −0.04), childhood maltreatment (g = −0.14, 95% CI [−0.44, 0.17]), past-year abuse severity (g = −0.04, 95% CI [−0.33, 0.27]), non-White race (OR = 0.73, 95% CI [0.37, 1.43]), and age (*g* = −0.08, 95% CI [−0.40, 0.21]).

To test whether representation in the imaging sample was especially low among women with the greatest psychiatric burden, we combined PHQ-9, GAD-7, and PCL-5 scores and compared the highest quartile with the remainder of the cohort. Seven of 52 women in the highest quartile had imaging data available (13.5%), compared with 54 of 155 women in the remaining three quartiles (34.8%, OR = 0.31, 95% CI [0.13, 0.71], *P* = .003). This difference was present before motion-based quality control.

To examine whether current safety context differed between the behavioral and imaging samples, we compared women reporting ongoing and past abuse. Five of 10 women reporting ongoing abuse had imaging data available (50.0%), compared with 47 of 121 women reporting only past abuse (38.8%, OR = 1.57, 95% CI [0.46, 5.40], *P* = .52), an estimate too imprecise to support a conclusion in either direction. Time since the most recent partner-inflicted BI was similar between groups (10.9 versus 10.9 years, g = 0.00, 95% CI [−0.36, 0.38], n = 37/85), although this measure was available for only 122 women.

### Motion quality control excluded women with the greatest BI burden

Participant-level head-motion quality control is standard practice in rsfMRI, namely as motion can produce spurious estimates of functional connectivity [43,44]. To quantify data loss at the quality-control stage, we applied the primary exclusion threshold of mean framewise displacement at or above 0.20 mm to the 61 women with rsfMRI data. The criterion retained 55 women (90.2%) in the analysis sample.

To test whether motion exclusion was concentrated among women with the greatest cumulative injury exposure, we compared women reporting ten or more partner-inflicted BIs with those reporting fewer BIs. Four of 11 women with ten or more BIs were excluded (36%), compared with two of 50 women with fewer BIs (4%, OR = 11.64, 95% CI [2.07, 65.45], *P* = .008) (Fig. 4). The median injury count was 11.5 among excluded women and 1.0 among retained women. The same pattern was present at alternative thresholds. At the least restrictive threshold of 0.30 mm, the only two excluded women reported ten or more BIs, while every woman reporting fewer BIs was retained (*P* = .030). At 0.15 mm, exclusion rates were 6/11 among women with ten or more BIs and 9/50 among women with fewer BIs (*P* = .019) (Supplementary Tables 3 and 4).

**Fig. 4.**
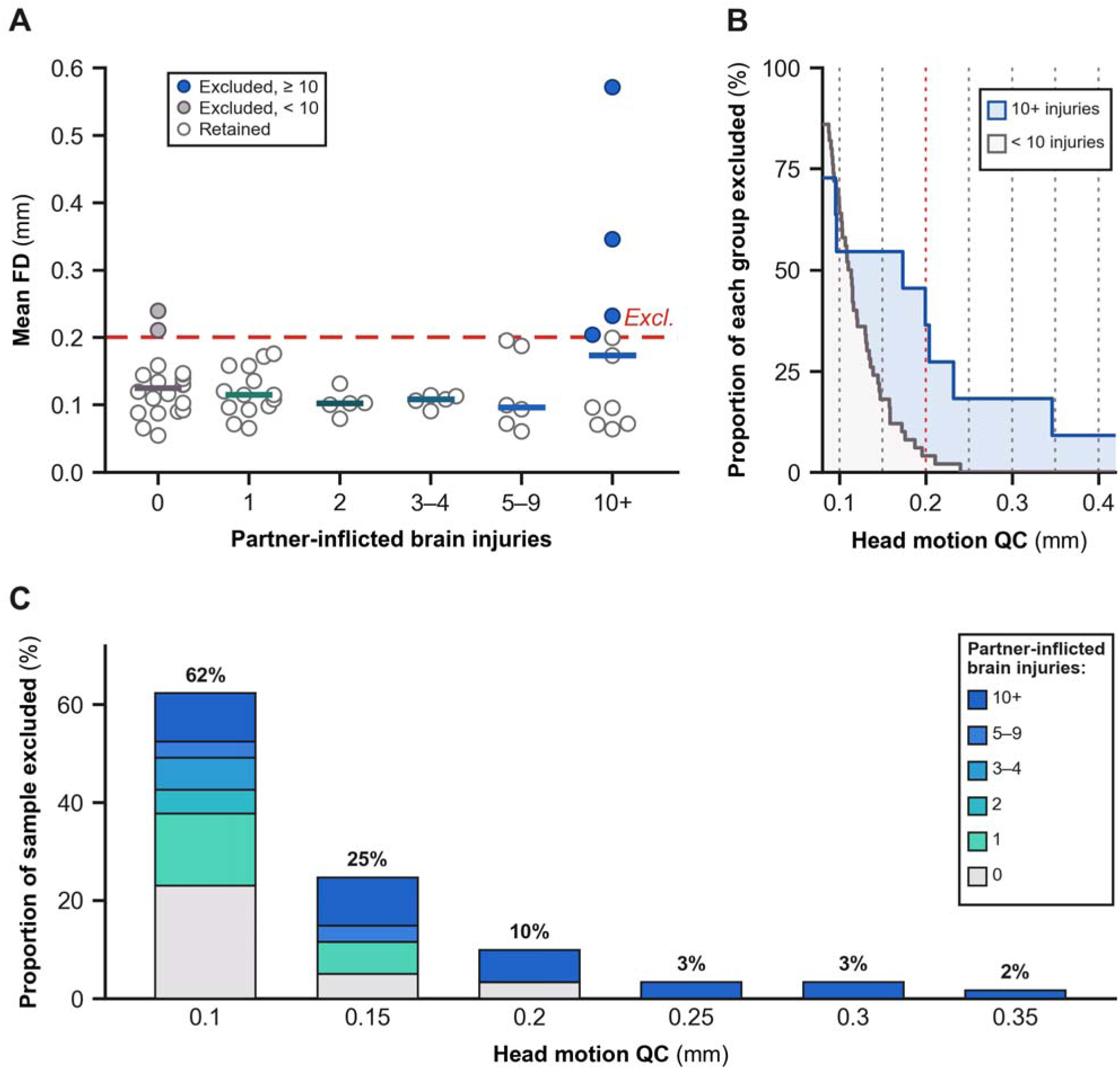
Representation bias in quality control of neuroimaging sample. **(A)** Mean framewise displacement for the 61 women with rsfMRI data, by cumulative partner-inflicted BI. Bars are group medians, colored as in **(C)**; the dashed line is the primary criterion of 0.20 mm. Open circles are retained women, gray circles excluded women with fewer than ten BIs, and blue circles excluded women with ten or more. Four of 11 women with ten or more BIs and two of 50 with fewer were excluded (two-sided Fisher exact *P* = .008, OR = 11.64, 95% CI [2.07, 65.45]). Head motion was unrelated to continuous injury count (Spearman ρ = 0.01, *P* = .918). **(B)** Percentage of each injury group that a criterion would remove, at every threshold from 0.08 to 0.42 mm; the red dotted line is the primary criterion. The difference between the groups is widest between 0.15 and 0.20 mm, the range in common use, and narrows at stricter and at more lenient thresholds. **(C)** Composition of the excluded set at six criteria, as a percentage of the 61 women, by injury band. Supplementary Table 5 reports leave-one-out analyses.

To determine whether the motion pattern extended across the injury distribution, we repeated the analysis using additional categorical and continuous injury measures. Exclusion rates were similar for women with any BI and women with none (4/41 versus 2/20, OR = 0.89, 95% CI [0.17, 4.59], *P* = 1.000). The difference was smaller at a threshold of five BIs (4/17 versus 2/44, *P* = .046) than at ten BIs (Supplementary Table 5). Mean framewise displacement was unrelated to continuous injury count (Spearman ρ = 0.01, *P* = .918, Pearson *r* = 0.26 for log-transformed injury count). The observed exclusions were therefore concentrated in the group with ten or more BIs rather than distributed monotonically across injury counts.

To assess whether the association depended on any one woman, we repeated the ten- or-more-BI comparison after removing each excluded participant in turn. This leave-one-out analysis tested whether a single observation accounted for the disproportionate exclusion of women with the greatest cumulative injury burden. Removing any one excluded participant with ten or more BIs changed the Fisher exact *P* value from .008 to .029 (Supplementary Table 5). No single excluded participant accounted for the direction of the association, although the wide confidence interval indicates that its magnitude requires confirmation in larger samples.

## Discussion

In this study, we investigated problems of representativeness and data exclusion at three major stages of IPV neuroimaging sample construction. At the eligibility stage, three in five women eligible for behavioral research were deemed ineligible for neuroimaging at initial screening. The most frequently recorded criteria were psychotropic medication use, geographic distance from the imaging center, and MRI contraindications. At the participation stage, enrolled women who underwent rsfMRI had significantly more education, were more likely to be employed, and reported lower psychiatric and neurobehavioral symptom burden than enrolled women without rsfMRI data. Finally, at the quality-control stage after rsfMRI, motion-based exclusion disproportionately removed women reporting ten or more partner-inflicted BIs. To our knowledge, this is the first empirical assessment of representativeness within IPV neuroimaging, and our findings empirically reveal that eligibility, participation, and quality control each bias the demographic, clinical, and BI burden represented in an rsfMRI sample.

Eligibility criteria substantially narrowed access to neuroimaging, and 23% of the women deemed ineligible failed only criteria that were later removed. This sensitivity to protocol design is important because IPV neuroimaging studies use heterogeneous experimental designs and eligibility criteria [49]. Scanner-safety restrictions determine who can undergo imaging, whereas clinical exclusions often reflect efforts to reduce sample heterogeneity. Geographic distance, our most common reason for those ultimately excluded, reflect the location of the imaging center rather than any characteristic of the participant. In IPV-related BI, psychiatric symptoms, medication use, prior injuries, and medical complexity may be features of the population under study rather than incidental sources of variation [50–52]. Excluding these characteristics can produce a narrower sample while changing the population represented and limiting comparison across protocols. Eligibility criteria should therefore be justified in relation to the scientific question and the population to which the findings are intended to generalize.

Participation favored women with more education and lower social and clinical burden. This pattern resembles population and clinical imaging studies in which participation varies with education, health, cognition, and clinical status [11–16]. The education difference is particularly consequential as IPV research increasingly examines associations between BI and cognition. Greater educational attainment may shift expected cognitive performance and restrict its observed range, which can obscure lower performance or impairment, and alter estimated brain–behavior associations. Related work in this cohort identified the greatest cognitive deficits among women reporting ten or more BIs even though the sample was more educated than the general population [34]. Demographic representation is especially important because the social conditions that vary across groups can shape both exposure to IPV and the neural and cognitive outcomes under study. Among trauma-exposed adults, neighborhood disadvantage has been associated with persistent neurocognitive deficits [53], greater threat-related neural activity [54], PTSD status, and perceived neighborhood danger [55]. Structural inequities have similarly been associated with racial and ethnic differences in neurophysiological tone after trauma [56]. Underrepresentation may therefore omit meaningful variation in the social conditions that contribute to cognitive, psychiatric, and neural outcomes.

Quality control selectively excluded women at the highest cumulative BI exposure, as every woman excluded at the 0.30 mm threshold reported ten or more BIs. Related work indicates that this subgroup has a distinctly lower cognitive profile relative to normative samples [34]. Removing these women narrows the ranges of both injury exposure and cognitive functioning, which can attenuate or obscure the brain–behavior associations most relevant to cumulative injury. Motion control remains necessary because motion can bias rsfMRI estimates [43,44]. Participant exclusion, however, is not analytically neutral when the excluded group differs systematically on the exposure or outcome under study [20–22,24,57,58]. The resulting estimates may be weaker, less precise, or otherwise distorted depending on how selection relates to the exposure and imaging outcome [59]. Quality-control decisions therefore require evaluation of both artifact reduction and changes in the population analyzed.

These analyses provide quantitative evidence for representational problems that IPV neuroimaging studies, including our own, often recognize qualitatively or assume. Statistical adjustment cannot recover women who never become eligible, do not participate, or are removed before analysis. Each source of selection should therefore be measured and addressed where it arises.

### Recommendations for representative neuroimaging of IPV

Given that selection can occur during eligibility, participation, and quality control, we ask: how can neuroimaging studies of IPV produce more representative samples?

At eligibility, distinguishing scanner-safety requirements from design exclusions would clarify the rationale for each criterion and show how defensible alternatives change the eligible population. Prospective records of screening counts, participant characteristics, exclusion reasons, and criterion changes would make these effects measurable. Comparisons between source and analytic samples can report effect estimates and uncertainty. These practices extend recommendations from alcohol-treatment and TBI research to examine how exclusion criteria affect generalizability [50–52]. They are especially important because IPV imaging cohorts remain modest, brain-wide associations often require far larger samples [60,61], and models may fail across racial and ethnic groups or for people who differ from sample stereotypes [58,59].

As a field, we would benefit from understanding why women from underrepresented demographic and clinical groups do or do not engage in research. IPV neuroimaging cohorts are difficult to recruit and remain small, so each participant affects both precision and sample composition. Research on how safety, trust, transportation, caregiving, symptoms, and perceived value shape participation could guide recruitment and compensation. Survivor and community partnerships can inform safe communication and acceptable procedures. Post hoc matching cannot replace this work because the matching fallacy can produce apparently comparable groups from the same unrepresentative population [62].

Importantly, access to research participation constrains the sample. Our recruitment reaches only women already connected to community partners, an online research platform, or social media, and geographic distance then narrows the group further. Indeed, among the 469 screened women who reported how they heard about the study, close to four in five named an online route, and fewer than one in twenty were referred by a shelter, domestic violence agency, or clinician. Of the 189 enrolled women whose residence at screening was recorded, only 2 were living in a shelter. Geographic distance was the most frequently recorded criterion still in use in our own cohort, and nearly 9% of women were deemed ineligible on this basis. Portable MRI may therefore bring protocols into communities that fixed-site imaging does not reach. A national survey found broad willingness among rural, Black, Hispanic, and economically disadvantaged respondents and identified trust and community engagement as central considerations [63].

At quality control, acquisition and analytic procedures can be evaluated for validity and representation. Shorter protocols, planned breaks, and behavioral preparation may reduce motion [64], while real-time monitoring can identify problems immediately when rescanning remains possible [65]. Future studies could compare censoring, denoising, and alternative inclusion approaches to determine which preserve valid inference with the least amount of selective data loss [66,67]. Similar recommendations have emerged in developmental neuroimaging and fMRI methodology, where quality-control pathways are evaluated for the samples they retain as well as the artifacts they reduce [24,58,67]. Reporting how each pathway changes sample composition would clarify the population represented by the final results.

Selection decisions determine which survivors inform estimates of IPV-related neural harm and how broadly those estimates can be interpreted. Consistent with Ricard et al. [25], transparent reports would describe sample composition and exclusions, show how acquisition and quality-control decisions changed representation, and limit interpretation to the retained population. Socially patterned exclusions or exclusions related to the exposure or outcome warrant cautious reporting of results and generalizability. Consistent reporting of race and ethnicity would also allow the field to measure progress toward representative datasets, while clearer journal standards could improve rigor and comparability [68]. These practices would make selection visible and allow readers to judge the claims that neuroimaging evidence can support.

### Representativeness as an ethical, legal, and societal concern

Addressing representativeness in the neuroimaging of IPV is especially pressing by nature of the population studied. In other fields, neuroimaging and related neuroscientific evidence already inform TBI diagnosis and rehabilitation planning [69], contribute to disability determinations [70], appear in legal proceedings [71], shape public policy [72], and have informed Supreme Court rulings on juvenile sentencing [73,74]. Findings from IPV neuroimaging are therefore likely to shape whether and how partner-inflicted BI is screened for, whether symptoms are recognized as possible consequences of injury, whether survivors receive rehabilitation, disability support, or compensation, how evidence is evaluated in the courtroom, and whether IPV-related BI receives public attention, policy responses, or legislative action. Yet our findings collectively suggest that the survivors most affected by IPV and partner-inflicted BI may also be the least represented in neuroimaging studies.

These ethical, legal, and societal implications make representativeness in IPV neuroimaging increasingly imperative. We are particularly concerned that unrepresentative samples may lead studies to understate the harms of IPV and partner-inflicted BI or fail to detect them altogether. Consequently, neuroimaging evidence intended to evaluate these harms could instead be used to minimize them in the courtroom, while modest or null findings could inadvertently discourage screening, recognition, or referral in clinical practice. The same findings could weaken the perceived need for services, research funding, and legislative action. Null findings from unrepresentative samples may also lead audiences, particularly those outside academic research, to mistake an absence of evidence for evidence of absence [75].

Finally, our concerns are heightened by neuroscience-related problems identified in the legal literature. The “seductive allure of neuroscience” describes how explanations and arguments, regardless of their merits, can become overwhelmingly more persuasive when accompanied by neuroscientific information such as brain scans [76]. In addition, the group-to-individual inference problem arises when a population-level finding is incorrectly assumed to apply to an individual case [77–79]. To illustrate, a study reporting no detectable cognitive harm from BI at the population level could be used to argue erroneously that an individual survivor did not experience cognitive impairment. Together, these ethical, legal, and societal problems underscore the risk that findings from unrepresentative samples may receive undue persuasive weight in high-stakes settings and be extended inappropriately to false conclusions about individual survivors.

### Limitations

Our study has several limits. As a cross-sectional study, we could not determine how participation and sample composition changed over time. Longitudinal work should therefore assess attrition and its effects on representativeness. Our data came from one cohort and imaging center, limiting generalizability. In addition, selection likely begins before the three stages we analyzed here, especially as recruitment through community partners or an online research platform reaches only women already in contact with those channels. Replication across cohorts and sites is needed to determine whether these patterns persist and whether sample composition differs across settings, particularly across international contexts. As reasons for nonparticipation in neuroimaging were not systematically recorded, we also could not determine why some imaging-eligible women who completed the behavioral study components did not undergo scanning.

Finally, other acquisition or quality-control methods may produce different analytic samples. Future studies should compare these approaches to determine which preserve valid inference with less selective data loss. We also lacked data to evaluate whether representation varied by sexual orientation or gender identity, although transgender populations experience disproportionate IPV [80]. Larger multisite studies should examine these factors and quantify how recruitment, portable imaging, weighting, and quality-control methods alter specific neuroimaging associations.

When our group published the first neuroimaging study of IPV and BI, our aim was to bring greater visibility to the harms of a widespread but often “invisible” trauma [6]. The present findings show that representativeness may be a necessary condition for achieving this aim, a concern now being raised across clinical neuroimaging more broadly [81]. We therefore encourage future researchers to rigorously examine who can enter, complete, and remain in an imaging study. As in doing so, we better ensure that neuroimaging evidence reflects the survivors whose harms we seek to make visible.

## Acknowledgments

We sincerely thank the women who took part in this study, and the study staff who contributed to data collection and curation. We also thank Harvard Catalyst for biostatistics consultation on the statistical analyses conducted in this study. CWM was supported by graduate funds from the Clarendon Fund, Harvard-UK Fellowship, and Knight-Hennessy Scholarship during research.

## Data and code availability

Direct identifiers have been removed from the analysis dataset. The de-identified data including the head motion estimates supporting this study are available from the corresponding author upon reasonable request. Raw data collected earlier in the study were uploaded to the Federal Interagency Traumatic Brain Injury Research (FITBIR) data repository at https://fitbir.nih.gov/, which is no longer open for further deposit.

## Supplementary Information

**Supplementary Table 1.** Women with resting-state fMRI data had more education, higher employment, and lower psychiatric burden than women without imaging data. To determine whether the imaging sample differed from the broader enrolled cohort, we compared 61 women with resting-state fMRI data and 146 without these data across socioeconomic, psychiatric, neurobehavioral, injury, abuse, and demographic measures. Values are means (SD) unless otherwise stated. Strongly right-skewed count variables are summarized with medians [IQR] and Cliff’s delta (δ), and the winsorized injury count is provided for comparison. Hedges *g* represents the standardized mean difference with percentile bootstrap 95% confidence intervals. Binary characteristics are summarized with Haldane-Anscombe-corrected odds ratios and 95% confidence intervals. The analyzed *n* is reported for each estimate and varies with missingness.

| Characteristic | Imaged | Not imaged | Effect size [95% CI] | $n$ |
| --- | --- | --- | --- | --- |
| Education (years) | 16.6 (2.6) | 15.4 (2.9) | 0.42 [0.13, 0.72] | 61/146 |
| Currently employed | 75.4% (46/61) | 59.3% (86/145) | OR 2.06 [1.06, 4.00] | 206 |
| Depression, PHQ-9 | 8.1 (5.2) | 10.2 (5.9) | -0.37 [-0.64, -0.09] | 61/146 |
| Anxiety, GAD-7 | 6.9 (5.2) | 8.9 (5.9) | -0.35 [-0.64, -0.07] | 61/146 |
| Post-traumatic stress, PCL-5 | 29.2 (17.4) | 35.3 (19.9) | -0.32 [-0.59, -0.02] | 61/146 |
| Neurobehavioral symptoms, NSI total | 20.3 (12.6) | 27.4 (14.9) | -0.50 [-0.78, -0.21] | 61/145 |
| NSI vestibular | 0.6 (0.6) | 0.9 (0.9) | -0.38 [-0.63, -0.11] | 61/145 |
| NSI somatosensory | 0.6 (0.5) | 0.9 (0.7) | -0.41 [-0.65, -0.15] | 61/145 |
| NSI cognitive | 0.9 (0.7) | 1.2 (0.8) | -0.32 [-0.61, -0.03] | 61/145 |
| NSI affective | 1.3 (0.8) | 1.7 (0.9) | -0.46 [-0.76, -0.17] | 61/145 |
| Ongoing abuse | 9.6% (5/52) | 6.3% (5/79) | OR 1.57 [0.46, 5.40] | 131 |
| Raw partner-inflicted BI count median [IQR] | 1 [0–5] | 1 [0–4] | $\delta$ 0.03 | 61/146 |
| Winsorized at 25 partner-inflicted BI count mean (SD) | 4.5 (7.1) | 4.2 (7.3) | 0.03 [-0.25, 0.34] | 61/146 |
| Strangulation-related BI count | 0 [0–0] | 0 [0–1] | $\delta$ -0.04 | 61/146 |
| Childhood maltreatment, CTQ | 56.2 (21.9) | 59.2 (22.1) | -0.14 [-0.44, 0.17] | 61/146 |
| Abuse severity, CASR-SF (past 12 months) | 6.5 (11.1) | 7.0 (11.1) | -0.04 [-0.33, 0.27] | 61/146 |
| Years since last IPV brain injury | 10.9 (9.0) | 10.9 (10.8) | 0.00 [-0.36, 0.38] | 37/85 |
| Non-White race | 25.0% (15/60) | 31.7% (46/145) | OR 0.73 [0.37, 1.43] | 205 |
| Age (years) | 40.2 (13.3) | 41.4 (13.1) | -0.08 [-0.40, 0.21] | 61/146 |
| Highest symptom-burden quartile | 11.5% (7/61) | 30.8% (45/146) | OR 0.31 [0.13, 0.71] | 207 |

**Supplementary Table 2.** Imaging criteria excluded three in five behaviorally eligible women. To identify the criteria that most often limited imaging eligibility, we summarized every recorded reason for neuroimaging ineligibility among 211 behaviorally eligible women. Women could meet more than one criterion, so percentages exceed 100% in total. The MRI contraindication item was unassessed in 58 women, producing different percentages for the full and assessed samples. Among ineligible women, 161 (76%) failed at least one criterion that remained in use, and 49 (23%) failed only criteria that were later removed. One participant was excluded for reasons noted during the interview, in which they discussed discomfort with MRI after a prior bad MRI experience.

| Criterion | *n* | % of excluded | % of those assessed | Assessed |
| --- | --- | --- | --- | --- |
| Criteria applied throughout |  |  |  |  |
| Geographic location | 64 | 30.3% | 33.7% | 190/211 |
| Magnetic-resonance contraindication | 58 | 27.5% | 37.9% | 153/211 |
| Schizophrenia, bipolar disorder or autism | 34 | 16.1% | 16.3% | 209/211 |
| Moderate or severe traumatic brain injury | 30 | 14.2% | 14.6% | 206/211 |
| Confounding neurological disorder | 29 | 13.7% | 14.4% | 201/211 |
| Substance use history | 28 | 13.3% | 13.6% | 206/211 |
| Brain injury within three months | 17 | 8.1% | 8.2% | 207/211 |
| Pregnancy | 1 | 0.5% | 0.5% | 197/211 |
| Prior bad MRI experience | 1 | – | – | – |
| Criteria since relaxed or dropped |  |  |  |  |
| Psychotropic-medication use | 105 | 49.8% | 51.5% | 204/211 |
| One or two partner-inflicted brain injuries | 58 | 27.5% | 29.1% | 199/211 |
| Age above the ceiling of 55 years | 38 | 18.0% | 18.5% | 205/211 |

**Supplementary Table 3.** Motion quality control disproportionately excluded women with ten or more partner-inflicted BIs across all evaluated mean FD thresholds. To determine whether this association depended on the mean FD threshold, we compared exclusion among women reporting ten or more BIs and women reporting fewer BIs at 0.15, 0.20, and 0.30 mm. Values are the number and percentage excluded within each injury group among 61 women with rsfMRI data. The 0.20 mm threshold was primary, and two-sided Fisher exact *P* values are reported.

| Threshold | Ten or more injuries excluded | Fewer than ten excluded | Fisher *P* |
| --- | --- | --- | --- |
| Mean FD ≥ 0.15 mm | 6/11 (55%) | 9/50 (18%) | .019 |
| Mean FD ≥ 0.20 mm | 4/11 (36%) | 2/50 (4%) | .008 |
| Mean FD ≥ 0.30 mm | 2/11 (18%) | 0/50 (0%) | .030 |

**Supplementary Table 4.** At the 0.30 mm mean FD threshold, the only women excluded reported ten or more partner-inflicted BIs. To describe how each quality-control threshold changed the excluded sample, we report the number and percentage excluded and the median BI count among excluded and retained women at 0.15, 0.20, and 0.30 mm. At the primary 0.20 mm threshold, the six excluded women reported 0, 0, 10, 13, 40, and 420 partner-inflicted BIs.

| Threshold | *n* excluded | % of sample | Median injuries, excluded | Median injuries, retained |
| --- | --- | --- | --- | --- |
| Mean FD ≥ 0.15 mm | 15 | 24.6% | 5.0 | 1.0 |

| Threshold | *n* excluded | % of sample | Median injuries,<br>excluded | Median injuries,<br>retained |
| --- | --- | --- | --- | --- |
| Mean FD ≥ 0.20 mm | 6 | 9.8% | 11.5 | 1.0 |
| Mean FD ≥ 0.30 mm | 2 | 3.3% | 11.5 | 1.0 |

**Supplementary Table 5.** The association between motion exclusion and ten or more BIs persisted after each excluded woman was removed in turn. To assess whether the result depended on the injury threshold or any one excluded observation, we repeated the primary 0.20 mm comparison at thresholds of any, five, and ten BIs and then removed each excluded woman in turn. Each row reports exclusion counts, an odds ratio with a 95% confidence interval, and a two-sided Fisher exact *P* value. Removing any one excluded woman with ten or more BIs changed P from .008 to .029. Mean FD was unrelated to continuous injury count, with Spearman ρ = 0.01, *P* = .918, and Pearson *r* = 0.26 for log-transformed injury count.

| Analysis | Exposed group<br>excluded | Comparison<br>excluded | Odds ratio [95%<br>CI] | Fisher *P* |
| --- | --- | --- | --- | --- |
| Ten or more vs fewer injuries | 4/11 (36%) | 2/50 (4%) | 11.64 [2.07, 65.45] | .008 |
| Five or more vs fewer injuries | 4/17 (24%) | 2/44 (5%) | 5.67 [1.07, 29.89] | .046 |
| Any injury vs none | 4/41 (10%) | 2/20 (10%) | 0.89 [0.17, 4.59] | 1.000 |
| Leave-one-out 1: excluded<br>participant with 0 injuries removed | 4/11 (36%) | 1/49 (2%) | 19.40 [2.62, 143.57] | .003 |
| Leave-one-out 2: excluded<br>participant with 0 injuries removed | 4/11 (36%) | 1/49 (2%) | 19.40 [2.62, 143.57] | .003 |
| Leave-one-out 3: excluded<br>participant with 40 injuries removed | 3/10 (30%) | 2/50 (4%) | 9.05 [1.50, 54.55] | .029 |
| Leave-one-out 4: excluded<br>participant with 420 injuries removed | 3/10 (30%) | 2/50 (4%) | 9.05 [1.50, 54.55] | .029 |
| Leave-one-out 5: excluded<br>participant with 10 injuries removed | 3/10 (30%) | 2/50 (4%) | 9.05 [1.50, 54.55] | .029 |
| Leave-one-out 6: excluded<br>participant with 13 injuries removed | 3/10 (30%) | 2/50 (4%) | 9.05 [1.50, 54.55] | .029 |

